# Long term recordings with immobile silicon probes in the mouse cortex

**DOI:** 10.1101/021691

**Authors:** Michael Okun, Matteo Carandini, Kenneth D. Harris

## Abstract

A key experimental approach in neuroscience involves measuring neuronal activity in behaving animals with extracellular chronic recordings. Such chronic recordings were initially made with single electrodes and tetrodes, and are now increasingly performed with high-density, high-count silicon probes. A common way to achieve long-term chronic recording is to attach the probes to microdrives that progressively advance them into the brain and isolate them from mechanical forces. Here we report, however, that such microdrives are not strictly necessary. Indeed, we obtained high-quality recordings in both head-fixed and freely moving mice for several months following the implantation of immobile chronic probes. Probes implanted into the primary visual cortex yielded well-isolated single units whose spike waveform and orientation tuning were highly reproducible over time. Although electrode drift was not completely absent, at least 70% of neurons retained their waveform across days. Thus, immobile silicon probes represent a straightforward and reliable technique to obtain stable, long-term population recordings in mice, and to follow the activity of populations of well-isolated neurons over multiple days.

## Introduction

An extraordinarily fruitful approach in systems neuroscience involves extracellular chronic recording of population spiking activity in awake animals. This technique has been especially useful in behaving rodents (Buzsaki, 2004; Stevenson and Kording, 2011), and is responsible for numerous advances in areas such as sensory-motor processing, navigation, emotion, and memory. It is likely to be employed even more widely in the near future, thanks to the forthcoming new generation of high-density, high-count silicon probes (Einevoll et al., 2012; Buzsaki et al., 2015).

Silicon probes provide an ideal combination of good isolation and large neuronal count (Csicsvari et al., 2003). They retain the excellent single-neuron isolation seen with traditional tetrodes (O’Keefe and Recce, 1993; Wilson and McNaughton, 1993), while providing a marked improvement in neuronal count. They are also arguably superior to bundles of microwires (Nicolelis et al., 1997; Freire et al., 2011; Vyazovskiy et al., 2011), as it is not clear that such (single) microwires can deliver multiple high-quality spike-sorted single units (Harris et al., 2000).

A common way to achieve long-term chronic recordings is to attach the probes to microdrives that progressively advance them into the brain and isolate them from mechanical forces. Microdrives have traditionally been used when implanting tetrodes (O’Keefe and Recce, 1993; Wilson and McNaughton, 1993). They were then modified for use with silicon probes, yielding successful recordings (Csicsvari et al., 2003). Microdrives allow to advance towards the target region in a gradual manner, which is thought to minimize the damage to the neural tissue, and to move past possibly damaged tissue, features that are considered to play an important role in the success of this approach (Buzsaki et al., 2015).

However, it is not known whether successful use of chronic silicon probes requires that they are moveable. Some of the first experiments involving silicon probes had them rigidly affixed to the skull (Hetke et al., 1994; Kipke et al., 2003; Moxon et al., 2004). However, since these initial reports, few laboratories have attempted chronic recordings with immobile probes, and success has been uneven. One study reported a failure to achieve recordings with reasonable signal to noise ratios (Karumbaiah et al., 2013). Others were more successful, e.g., (Retailleau et al., 2013; Kozai et al., 2015a), but did not provide quantitative measures of the probes’ ability to record well-isolated neurons with stable waveforms. Specifically, none of the studies provided evidence that the same neurons can be recorded over multiple days.

Here we report that it is possible to obtain high quality spiking recordings in mouse cortex for several months following the implantation of immobile chronic probes. Immobile probes implanted in primary visual cortex yielded well-isolated single units with highly reproducible visual responses. Using these visual responses, we estimate that well over 70% of the units remain stable across days. These results indicate that long-term chronic recordings can be performed with no microdrive, in both head-fixed and freely moving mice, with most of the recorded units remaining stable across days.

## Methods

All experimental procedures were conducted according to the *UK Animals Scientific Procedures Act 1986 Amendment Regulations 2012*. Experiments were performed at University College London under personal and project licenses released by the Home Office following institutional ethics review.

### Head plate implantation

Mice of either sex were implanted with a custom-built head plate (stainless steel, 1.1 g; Fig. 1). The animals were anesthetized with isoflurane, and kept on a feedback-controlled heating pad (ATC2000, World Precision Instruments, Inc.). Hair overlying the skull was shaved. Next, the skin and the muscles over the central part of the skull (approximately between the bregma and lambda) were removed. The skull was thoroughly washed with saline, followed by cleaning with a 3% solution of hydrogen peroxide. The head plate (Fig. 1) was held in its intended position by an alligator clip, while dental cement (Super-Bond C&B; Sun Medical, Japan) was applied to attach it to the skull. The exposed part of the skull within the hole in the head plate was covered with Kwik-Cast (World Precision Instruments, Inc.). The entire procedure was typically completed in < 40 minutes. Post-operative pain was prevented by administering a non-steroidal anti-inflammatory agent (Rimadyl) before the procedure and on the three following days.

**Figure 1.**
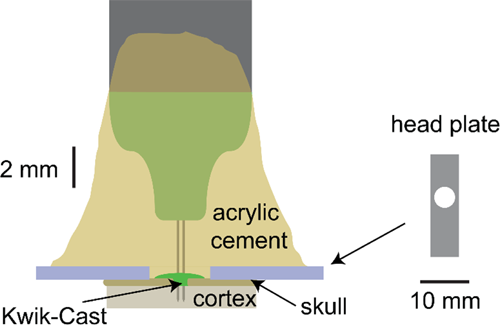
The chronic implant. *Schematic of the implantation (for 2-shank CM16 NeuroNexus probe) and the head plate. The total weight of the head plate, 16-channel probe and the cement used for attachment to the skull is ∼2 gr (2.5 gr if a bigger 32-channel probe is used).*

### Probe implantation

In principle, the probe can be implanted right after the head plate. However, it can be advantageous to postpone its implantation, so that the animal can first be trained in a behavioral task that requires head fixation. Moreover, probe implantation might be best carried out in a separate rig dedicated to electrophysiology. We thus implanted the probe days or weeks after the animals had recovered from the head plate implantation.

For probe implantations, mice were again anesthetized with isoflurane and kept on a heating pad. We removed the Kwik-Cast skull cover and drilled a small incision in the cranium (burr #19007-07, Fine Science Tools). Following this craniectomy the brain was protected with Ringer solution. We then lowered the probe through the dura using a manipulator (PatchStar, Scientifica, UK) to a required depth. For primary visual cortex recordings (3.5 mm posterior and 2.5 mm lateral to bregma), we aimed for a depth of ∼500-750 μm below the cortical surface, to reach the deep layers. In all but one case (see below), we used a CM16 or CM32 probe with 1-4 shanks from NeuroNexus, Inc; in the majority of the animals these were 2-shank CM16 probes, with 2 groups of 4 nearby sites (i.e., two tetrodes) on each shank (NeuroNexus A2x2-tet-3mm-150-150-121).

Once the probe was in its required position, we soaked up most of the Ringer solution and covered the craniectomy with a thin layer of Kwik-Cast. Finally, we applied several layers of acrylic cement (Simplex Rapid, Kemdent, UK). Each layer of the cement was allowed to fully cure before the next one was applied. For the first layer, the cement should be liquid enough to flow around the probe’s insertion site, and during its application one needs to be careful not to touch the probe directly. The cement for the second and subsequent layers can be more viscous, resulting in shorter curing time. This procedure firmly affixed the probe to the cranial cap formed by the Super-Bond C&B cement and the head plate.

The hardening of the acrylic cement around the probe causes displacements of ∼10-20 µm (Lee et al., 2014), which did not appear to pose a problem for extracellular silicon probes. A schematic, showing to scale an implanted probe (CM16 probe with 2 shanks) is presented in Fig. 1. The same approach can be used to implant more than one probe in an animal, as we have successfully done in one of the mice.

During the entire implantation procedure the probe was connected to an amplifier, to monitor LFP and spiking activity. This helps to position the probe at the desired depth and location (e.g., to target a particular retinotopic position in area V1), whereas abrupt changes or disappearance of neural activity are a clear sign of problems with the implantation. To increase signal quality, prior to implantation we electroplated the probes with the polymer PEDOT:PSS (Ludwig et al., 2006), using an appropriate device (nanoZ, Multi Channel Systems, Germany). Once the final layer of acrylic had cured, the amplifier head-stage was detached from the probe by carefully pulling it up.

We kept the animals anesthetized during the entire procedure (which lasted 1.5-2.5 hours), but anesthesia is strictly necessary only for the craniectomy. After the craniectomy, one could let the animal recover in its home cage, and perform the implantation later, while the animal is head-fixed and awake. Indeed, probe insertion causes no distress to the animal. Similarly, the cementing procedure is non-invasive: Kwik-Cast prevents any direct contact between acrylic cement and live tissue. This possibility may be preferred if spiking activity needs to be observed during wakefulness at the time of implantation.

The basic approach described above assumes that the silicon probe is rigidly attached to the head-stage connector. In one pilot experiment we used similar techniques to implant a tethered probe (NeuroNexus H32). We attached the probe to a narrow band of duct-tape which in turn was held by an alligator clip controlled by the PatchStar manipulator. The silicon probe was inserted into the cortex and cemented in its final position. After the cement holding the probe had cured, the duct-tape was cut, and the head-stage connector along with the tether cable were cemented to the skull.

A further refinement of these methods may include the administration of dexamethasone to reduce potential inflammatory responses to the probe, e.g., (Polikov et al., 2005; Kipke et al., 2008; Vyazovskiy et al., 2011).

### Electrophysiology and spike sorting

Recordings were performed using the Cerebus (Blackrock Microsystems, Salt Lake City, Utah) or the OpenEphys (Siegle et al., 2015) systems. Broadband activity was sampled at 30 kHz (band pass filtered between 1 Hz and 7.5 kHz by the amplifier), and stored for offline analysis. For reference signal we used either the head plate or a small golden pin implanted over frontal cortex during the original surgery.

Spikes were sorted using the KlustaSuite software (Rossant et al., 2015), freely available on the Web (klusta-team.github.io). This suite involves three software packages, one for each of three steps: spike detection and extraction (SpikeDetekt), automatic spike clustering (KlustaKwik), and manual verification and refining of the clusters (KlustaViewa). The first step involved detecting spike waveforms in high-pass filtered traces, as in the older NDmanager software suite for tetrode recordings (Hazan et al., 2006). For the second step, we extracted 3 principal components from the spike waveform on each channel, and used them in an automated clustering analysis (Kadir et al., 2014). The third step involved visual inspection of the spike waveforms, and auto- and cross-correlograms of the clusters (Hill et al., 2011). If this inspection revealed that the cluster contained spikes from more than a single neuron (“under-clustering”), the cluster was manually split. Conversely, if several clusters contained spikes of the same single neuron (“over-clustering”, which is most often revealed by inspection of the auto- and cross-correlograms of the clusters) they were merged. In addition, clusters without a clean refractory interval were discarded (Rossant et al., 2015). All the steps were oblivious to sensory responses of the units. For experiments done on several consecutive days, the recordings were concatenated, and the spike sorting was performed as if dealing with single, continuous recording. Time was not used as one of the dimensions for cluster separation.

### Visual stimuli and data analysis

On 4 consecutive days we presented mice with 60 randomized sequences of a set of 8 drifting gratings differing in orientation and direction (2 Hz, 0.04 cycles/degree, 100% contrast, 1 s stimulus duration and >2 s inter-trial interval). In these experiments, across the 4 days the animals were head-fixed in the same location with respect to the screens on which the stimuli were presented. The visual stimuli were presented on two of the three available LCD monitors, positioned 25 cm from the animal and covering a field of view of 120 x 60 deg.

Peri-stimulus time histograms (PSTHs) were computed using 1 ms bins over a 2 s interval (1 s for the stimulus and 500 ms pre and post stimulus, to include baseline activity and possible offset responses) by averaging across trials and smoothing with a 15 ms boxcar window. For every unit the presence of robust sensory response on each day was assessed by correlating the PSTHs computed from the first and last 30 trials. A unit was considered to have a reliable sensory response on a particular day only if this correlation exceeded 0.3 for at least one of the eight stimuli.

### Cluster stability estimate

Because spike sorting was performed blind to the sensory responses of the neurons, the stability of neuronal sensory responses could be used to derive an estimate of the fraction of clusters corresponding to stably-recorded neurons.

The method starts by counting the number of neurons with stable sensory responses across consecutive days. Specifically, for every cluster *c* with reliable visual responses on days D and D+1, we compute the correlation of cluster *c*’s visual response on day D+1 to the responses on day D of all the clusters with reliable visual responses on the same tetrode (we denote the number of such comparisons as *L*). If cluster *c*’s response on day D+1 was more similar to its own response on day D than to any of the other *L* – 1 clusters’ responses on day D, this is denoted a “match.”

We estimate the confidence interval using a probabilistic model of cluster stability. Let *s* denote the probability that a cluster is truly stable across the two days; our aim is to estimate *s*, although we cannot directly observe it. Let q denote the probability that a match will be found for a given cluster, if that cluster represents a stable neuron (this probability should be close to 1, but may be smaller in the case another neuron on the tetrode has a similar sensory response). Let *r* denote the probability that a match will be found if the cluster does not represent a stable neuron, but represents different cells on the two days; by a symmetry argument, this probability will be at most 1/*L*. We assume that the match probability is independent between neurons, conditional on their stability.

Let *m* denote the (marginal) probability of observing a match. By the formula of total probability we have *m* = *qs* + *r*(1 – *s*) = *r* + *s*(*q* – *r*).

Using the fact that *q* ≤ 1 and 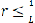, we can show that

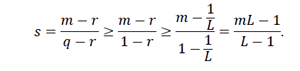

Our strategy is as follows. By observing the total number of matches found in the data, we obtain a (one-sided) binomial confidence interval for *m*, i.e. a value *m*_0_ such that *m* > *m*_0_ with 95% confidence. We then apply the above inequality to obtain from this a lower bound for *s* that holds with 95% confidence.

## Results

We implanted 11 mice with immobile, chronic 16- or 32- contact NeuroNexus probes (CM16 and CM32 configuration; Fig. 1), and one mouse with a similar probe but with a flexible connector (H32, NeuroNexus). The probes were located in the primary visual cortex (V1), or in the frontal cortex. Recordings began several days after the implantation, and were generally carried out in awake head-fixed animals, as this allowed us to precisely control the visual input.

We routinely obtained high quality recordings several months after probe implantation (Fig. 2). Of 11 animals implanted with fixed CM-type chronic electrodes, 9 provided high-quality recordings as long as recordings continued; in the remaining two, most spiking activity disappeared after about one week, and recordings were discontinued. In chronic recordings from the other 9 animals the incidence of large spikes ( > 80 μV) on the best site of each probe resembled data obtained with acute silicon probes (Fig. 2D), suggesting that any mechanical and inflammatory damage caused by the immobile chronic probe in these mice was not sufficiently severe to disrupt what appears to be standard electrophysiological activity in the nearby cortical tissue.

**Figure 2.**
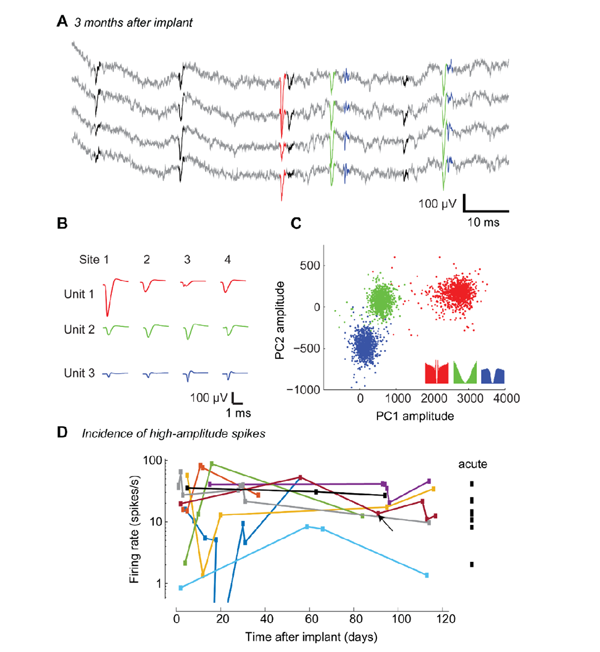
Recording with chronically implanted, immobile silicon probes. **A.** Raw traces from four sites arranged in a tetrode configuration, recorded 91 days after implantation of the silicon probe (A2x2-tet-3mm-150-150-121). Spikes detected as threshold crossings after high-pass filtering are shown in black or color. **B.** Average spike waveforms of three units shown in A (red, green, and blue), for each tetrode contact. **C.** 1,000 randomly chosen spike waveforms of each unit, projected on 2 out of the 3x4=12 dimensions of the space of principal components used for clustering. **Inset:** autocorrelograms for the three units, during spontaneous activity. **D.** Recording quality in 9 mice with chronic implants, assessed by the rate of spikes with trough amplitude exceeding 80 μV for the best site. For comparison, the same analysis in 8 acute recordings is shown on the right. The example recording shown in panels A-C is indicated by an arrow.

These data demonstrate that high quality units can be recorded for several months after implantation of fixed silicon probes. We next asked an additional question: how frequently might an experimenter expect to record from at least some of the same neuronal population over multiple days? In rats, for experiments performed with silicon probes using microdrives, it was reported that recordings from the same units occasionally last across days, e.g., (Diba et al., 2014; Schwindel et al., 2014). We expected the position of probes affixed to the skull to be no less, and likely even more stable than with a microdrive. Thus we proceeded to investigate the stability of the recordings across days in a systematic and quantitative manner, using the consistency of the neurons’ sensory responses to obtain a lower bound for the fraction of units that are recorded stably.

Our aim in these next analyses is to estimate the fraction of stable neurons, not to produce a method that indicates which clusters are stable in a given recording. Indeed, a metric based on receptive field similarity would be unsuitable for such a purpose. For example, using such a metric in studies of plasticity would lead to biased results (an underestimate of receptive field changes). To determine which recorded units are stable, therefore, a method based solely on waveform shapes should be used, such as those proposed by Tolias et al. (2007), or Eleryan et al. (2014).

To estimate the fraction of stable recorded units, we analyzed data from two mice in our sample, who had been presented with drifting gratings (8 directions) for 4 consecutive days (stimuli suitable for stability analysis had not been presented to the other mice). For these data, spike sorting was performed for the entire four days of recording together, so that the procedure was oblivious to the day in which each spike occurred as well as the neuron’s sensory tuning, and assigned spikes to clusters only based on spike waveform. Manual curation of these clusters was performed by an operator without access to the sensory tuning.

Many of the resulting clusters had stable visual response properties for multiple days. Consider four example units recorded on the same tetrode during four consecutive days (Fig. 3A). Although these neurons were close neighbors, they differed in the dynamics of spontaneous and evoked firing rates, in temporal profile of the response, and direction selectivity (Fig. 3B). This is consistent with the salt-and-pepper organization of rodent V1 and the absence of orientation columns (Ohki et al., 2005). While the measured peak response of the neurons occasionally fluctuated, the orientation tuning and temporal profile of these neurons’ responses was stable across the four recording days (Fig. 3B).

**Figure 3.**
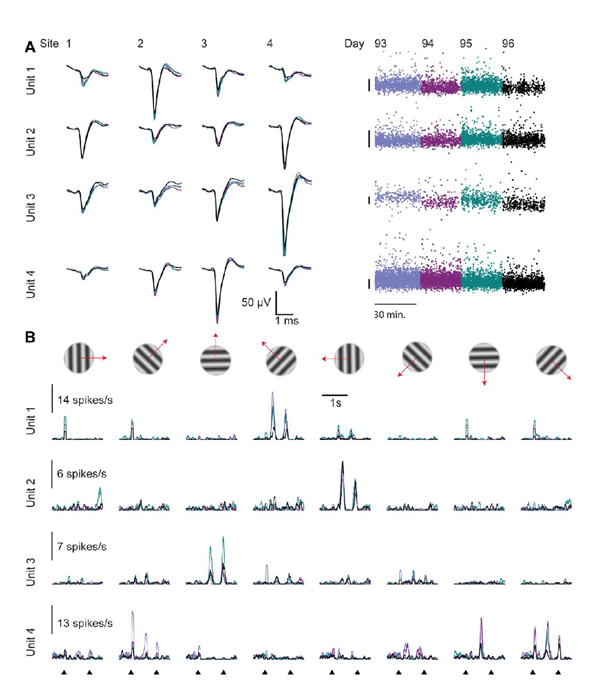
Examples of stable visual responses across days. Four units recorded on one tetrode for four consecutive days. **A. Left:** Mean waveforms of the example units (each day in different color, overlaid). **Right:** The amplitude of each spike (on the site with the largest waveform). Scale bars: 100 μV. **B.** Responses of the four example units to drifting gratings (8 directions), on four consecutive recording days (60 trials per direction per day). Onset and offset times of the gratings are indicated at the bottom.

Other units, however, showed evidence of drift in spike waveform and response properties when recorded across days (Fig. 4). For instance, the first example neuron shown in Fig. 4 seemed to disappear on the 4th day. Such occasional disappearances are not likely to be caused by the death of the neurons, because these recordings were performed weeks or even months after the implantation, when inflammatory responses are no longer likely to have a major day-to-day impact. Furthermore, we also observed units that appeared after the first day. For instance, the second unit in Fig. 4 seemed to appear on day 2: its firing rate on day 1 was 7 times lower than on the following days. A likely technical explanation for appearances and disappearances is mechanical displacement of the probe with respect to the neuron. Because of such drifts, on some days the spikes of a neuron would stand out of the multiunit activity as a well isolated cluster, while on other days they blend into it. This said, bona-fide changes in the neurons’ firing rate are also possible, and so is a change in spike waveform as a result of morphological plasticity or altered ion channel expression.

**Figure 4.**
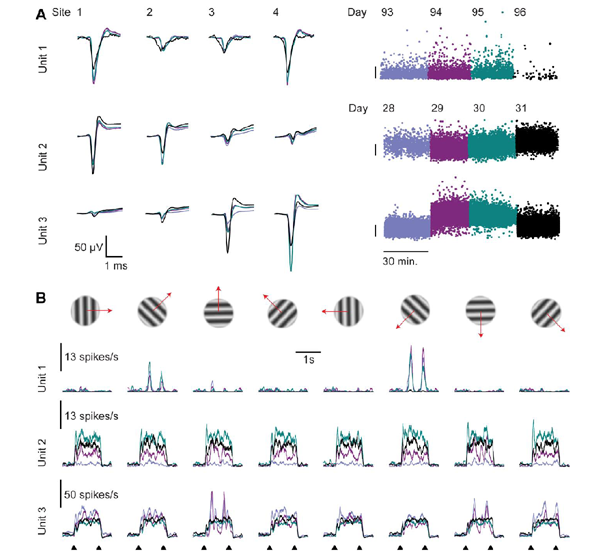
Example of unstable units across a 4 day recording. Three example units whose recorded responses changed markedly across days. Unit 1 is from a different shank of the recording shown in Fig. 3. Units 2, 3 were recorded on the same shank in the second animal. **A. Left:** Mean waveforms of the example units (each day in different color, overlaid). **Right:** The amplitude of each spike (on the site with the largest waveform). Scale bars: 100 μV. **B.** Responses of the example units to drifting gratings (8 directions), on four consecutive recording days (60 trials per direction per day). Onset and offset times of the gratings are indicated at the bottom. The example units illustrate three possible scenarios of drift. Unit 1 disappeared on the last day (and the few spikes of similar waveform detected on this last day probably do not originate from the same neuron). Unit 2 appears on the 2^nd^ day – its firing rate on day 1 was 7 times lower than on the following days. Unit 3 exemplifies a complex drift pattern where both the waveform and the response profile change across days.

To statistically analyze the stability of the recorded units, we began by comparing the response variability of the same putative unit across days to the variability between units (Fig. 5). We restricted the analysis to units with reliable temporal response profile within individual sessions on a single day, which was defined as correlation of at least 0.3 between PSTHs computed from two halves of the repeats, for at least one of the eight directions (repeating the analysis with other thresholds in the 0.2-0.5 range provided quantitatively similar results, data not shown). Overall, we recorded from 46 such units for more than one day. Of these, 23 units were tracked for all four days, 9 units were tracked for three days and 14 units were tracked for only two consecutive days (thus allowing for a total of 23 · 3 + 9 · 2 + 14 = 101 consecutive day comparisons). In addition, 18 visually responsive units were encountered only on one of the four days. Of the combined 64 units, 14 were narrow spiking (putative interneurons) and the rest were wide spiking, as judged by their mean spike waveform (Barthó et al., 2004). The measured peak to peak mean spike amplitude was 116 ± 26 μV (median ± MAD).

**Figure 5.**
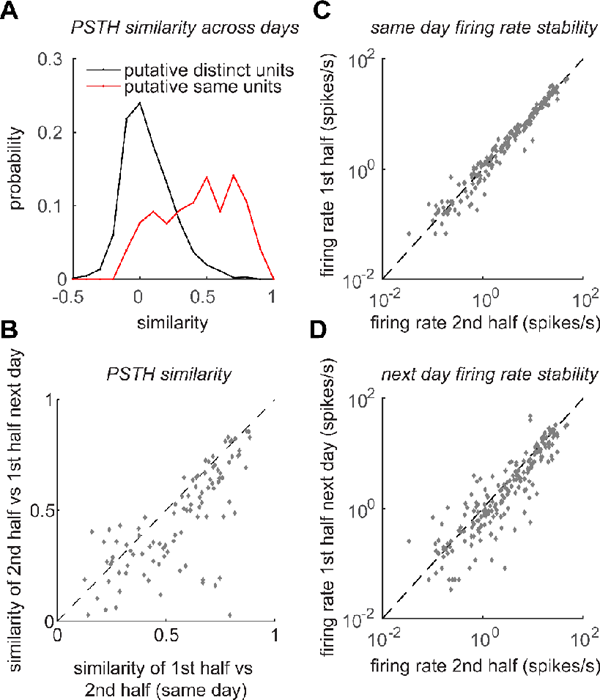
Stability of recordings across days. **A.** Distribution of signal correlations across days between putative distinct units (black) and between units that were judged to be the same by semi-automatic spike sorting (red). The two distributions are markedly different (P<10^−100^, ranksum test). **B.** Similarity (correlation) of response temporal profiles computed separately from first and second halves of trials on the same day vs similarity of responses computed from similar number of trials from two subsequent days (*ρ* = 0.75, P<10^−19^, 101 data points from 46 units in 2 animals). **C.** Firing rate in first vs second half of trials on the same day (ρ = 0.98, P<10^−100^). **D.** Comparison of firing rates in second half of trials and first half of trials on the following day (ρ = 0.82, P<10^−50^). In C,D only the preferred direction of each unit and a direction orthogonal to the preferred were used (202 data points from 46 units).

For units that were tracked for more than one day, we computed the correlation between PSTHs of individual units on subsequent days (8 PSTHs per unit per day) and compared them to correlation of PSTHs of pairs of units recorded (on the same tetrode) one day apart. Under the null hypothesis of no relationship between units recorded on subsequent days, these two sets of correlations are indistinguishable. However, this null hypothesis was clearly incorrect (Fig. 5A). Whereas the latter correlations were distributed around 0, just as one might have expected (an average zero signal correlation), the former correlations were positive, and the two distributions were markedly distinct (P<10^−100^, ranksum test).

To further assess the stability of the recording, we compared the correlation between PSTHs computed from the first and second halves of the trials on a given day and across days. We represented the responses of each neuron on any given day by two vectors of 16 · 103 entries each (one vector for responses in the first and second halves of the trials, correspondingly), these vectors included the concatenated PSTHs of the responses to each direction. We found the correlation of similarity within days and across days to be high (ρ = 0.75) and highly significant (P<10^−50^). In the majority of the cases, the time course of the response was about as reliable across consecutive days as it was across a single recording session, as shown in Fig. 5B. The few points in Fig. 5B that are furthest from the diagonal represent units that had a profound change in their spike waveform (a difference of 30-50% in the amplitude of the mean waveform) or in their firing rate (a difference of 1000-2000%) or both, which suggests that the classification of these as a single unit might have been wrong as a result of a drift, like in the examples in Fig. 4. For the majority of the cases, however, the firing rate was highly stable (ρ = 0.82, P<10^−100^, Fig. 5C,D). One also needs to keep in mind that we made no attempt to match brain state and animal behavior across sessions, and these factors contribute to fluctuations in firing rate (Reimer et al., 2014; Scholvinck et al., 2015; Vinck et al., 2015).

While the analyses in Fig. 5 allow to reject the null hypothesis of absolutely no stability, and provide a qualitative picture indicating that stability is the rule rather than the exception, they do not provide a quantitative measure of how common stable units are. To address this point, we devised a way to estimate a 95% confidence bound for the fraction of stable recorded units, by comparing the visual response of a given cluster on one day to the visual responses of itself and other clusters on the same tetrode on the preceding day (see Methods). Intuitively, the idea is that if cluster *c* indeed represents the same neuron on both days, the visual response on the preceding day most similar to *c*’s response on the second of the two days would belong to *c* with high probability, resulting in a ‘match’. If, on the other hand, this cluster corresponds to two different neurons on the two days, the probability of a match is inversely proportional to the number of clusters we compare against.

With the collected data, we had 93 cases where a response could be compared against at least 4 other units on the same tetrode on the preceding day. Of these, 78 were matches, and 15 were mismatches. Using the inequality derived in the Methods section (and substituting L = 5), we find a 95% confidence lower bound for the probability of a match to be at least 0.76, and hence a 95% confidence lower bound for the probability of a unit being stable, of 0.7. For 99% and 99.9% levels of confidence, the lower bounds on the probability of a stable unit are 0.66 and 0.62, correspondingly.

The estimate of 70% stability that we get from this analysis is likely to be an underestimate of the number of stable units, for the following three reasons. First, 70% is a lower bound, i.e., the value that can be rejected at the 0.05 confidence level, and the actual value is likely to be higher. Second, the analysis uses the conservative assumption that for a stable unit a match happens with probability of 1, which is not the case in practice, because nearby units can have highly similar responses (e.g. see Fig. 4). Third, most of the matches were on tetrodes where more than 4 additional visually responsive units were recorded, meaning that the probability of a match for an unstable unit was even less than 1/5 (the value used above).

One final piece of evidence in favor of concluding that stability is a ubiquitous feature in our recordings comes from several occasions where we observed cross-correlograms of the rare putative monosynaptic connections, and these were also conserved across days or even weeks, as shown in an example in Fig. 6. The similar shape of spike waveforms (Fig. 6A and C), and of their autocorrelograms (inset in Fig. 2C) and crosscorrelograms (Fig. 6B and D), is consistent with the very same 3 neurons being recorded. Most tellingly, the cross-correlograms between neurons 1 and 3 and between neurons 2 and 3 contain a sharp peak right after time zero, a pattern strongly suggestive of monosynaptic connection between these neurons. Because such patterns are rarely observed in cortex (Barthó et al., 2004), the most plausible explanation is that the very same triplet of neurons was recorded 20 days apart.

**Figure 6.**
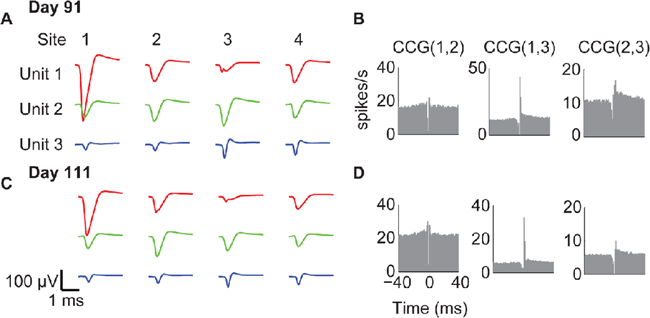
Stability of spike waveforms and spike train statistics across recording sessions. **A.** Spike waveforms of three units, for each contact of the tetrode (columns), same units shown in fig. 2B-C. **B.** Crosscorrelograms (CCGs) for the pairs of these three units, suggesting a monosynaptic connection between units 1,2 and unit 3. **C-D.** As in (A-B), for a recording performed 3 weeks later. The similar structure of spike waveforms, autocorrelograms (not shown) and crosscorrelograms, and particularly of crosscorrelograms of putative monosynaptically connected pairs, suggests that the very same 3 neurons were recorded on both occasions.

## Discussion

Our results demonstrate that recording spiking activity with immobile silicon probes in mouse cortex is feasible and straightforward, and provides high-quality recordings of well isolated units for at least several months after the probe implantation: we obtained successful recordings with this method 9 times out of 11. Furthermore, the recordings from individual units can be stable across periods of several days and above.

Compared to recordings with probes mounted on a microdrive, our approach offers multiple advantages. First and foremost, the implantation procedure is considerably briefer and simpler, reducing the chance of mistakes. Second, without a microdrive the implant needs less “real estate” on the skull and is potentially lighter. These advantages can allow implanting probes simultaneously into multiple areas of the brain, in configurations that would have been complicated or impossible with microdrives. Third, it is likely that the recordings in the proposed configuration are more stable than with a microdrive, as there are no moving parts.

Of course, these advantages come at a price of not being able to move the probe. Slowly advancing the probe into the brain area of interest is thought (albeit not proven) to cause less damage than direct implantation (Buzsaki et al., 2015). In addition, the microdrive allows one to move further if the recording quality deteriorates, or to reach a different brain region. An additional disadvantage of the technique in its present form, when compared to some ways of performing a microdrive implantation, is that the probes cannot be recovered.

The use of probes rigidly affixed to the skull may also become more important with the introduction of a new generation of high-count, high-density silicon probes. These probes will contain around 1,000 contacts spread over a length of ∼1 cm (Buzsaki et al., 2015), which is from most angles longer than the entire mouse brain, and spans the entire travel range of typical microdrives. Such probes might allow one to follow individual units as they shift coherently across drift events, which is typically not possible with the current probes. An additional possible application of the method is in combination with a chronic window for imaging, for a combined electrophysiological and 2-photon imaging interrogation of the same population of neurons (Kozai et al., 2015b).

As in other approaches to chronic recordings (Polikov et al., 2005; Kipke et al., 2008; Buzsaki et al., 2015), in our results the number of well-isolated units and their stability varied not only across animals but also for different shanks and tetrodes in the same animal. For example, in the recording shown in Fig. 3, three of the tetrodes yielded 5-12 good units each, whereas the fourth tetrode yielded only one well isolated unit. Yet, this “underperforming” tetrode was just 150 μm away from two “well performing” tetrodes, and on the same shank as one of them. One possible reason for such disparities is a local inflammatory response of the cortical tissue to the implanted probe (Polikov et al., 2005; Potter-Baker et al., 2014; Jorfi et al., 2015).

So far, few other laboratories have attempted chronic recordings with immobile probes. Reports that pioneered the use of immobile silicon probes were published over a decade ago (Hetke et al., 1994; Kipke et al., 2003; Moxon et al., 2004); yet, their lead was followed only by a few studies, with uneven success. At the time of writing, NeuroNexus CM-type silicon probes are the most ubiquitously available models for recordings without a microdrive, but a literature search (Google Scholar) for the terms “NeuroNexus” and either “CM16” or “CM16LP” or “CM32”, found only three such works in rodents. Karumbaiah et al. (2013) report a negative result, i.e., a failure to achieve recordings with reasonable signal to noise ratios using NeuroNexus CM32 and H32 probes, in contrast to recordings with floating microwires. Retailleau et al. (2013), however, used CM16 probes to record from freely moving rats, and obtained good recordings, stable over consecutive days (at least as judged by mean firing rate). Similarly, Kozai et al. (2015a) followed a procedure similar to the one described here (except for the use of head plate) to implant linear 16-contact probes into mouse V1. They then recorded for several months to investigate multiunit and LFP activity in different layers. Their conclusions are comparable with ours, supporting the feasibility of using immobile silicon probes for chronic recordings in mice.

The reasons for the success of our methodology, compared to other approaches to fixed probe recordings (e.g. Karumbaiah et al. (2013)) are not clear. Perhaps an important reason for the success of our approach is the use of a head plate as part of the implant (Fig. 1). The design of present-day interconnects between probe and amplifier head-stage is such that a significant force has to be applied for connecting the two. This is a major source of mechanical instability. The head plate is likely to mitigate the problem by reinforcing and protecting the skull around the implantation site. In addition, we always plugged in the head-stage when the animal was head-fixed (releasing it into its home cage afterwards for freely moving recordings). Thus the pressure was transferred to the head plate and the head-fixation apparatus, and never applied directly to the skull. However head-fixation during the actual recording is not a requirement. Indeed we have performed several test recordings in an animal walking and sleeping in its home cage, which also produced good results (not analyzed here).

To quantify recording stability, we introduced a method that relies on the unique sensory response profiles of the neurons. This method resembles the one used in a study in primate IT cortex, where the unique fingerprint of IT neurons was formed by responses to hundreds of images (McMahon et al., 2014). In mouse V1 neurons, we found that responses to drifting gratings are sufficient, as the PSTHs of even neighboring neurons exhibit a large diversity. Using visual responses as an independent measure for stability, we were able to reject the null hypothesis that less than 70% of the units considered stable across days by the spike-sorting procedure correspond to single neurons. While 70% is a bound that we were able to reject as too low at the P=0.05 level of confidence, the actual probability is likely to be 10%-20% higher, and can be further improved by not considering units that undergo a profound change in spike size and waveform, or firing rate.

Several recent studies in macaques also report the ability to achieve stable recordings for days or even months from the same units using microwires (Jackson and Fetz, 2007; Tolias et al., 2007; Hall et al., 2014; McMahon et al., 2014) or Utah arrays (Dickey et al., 2009; Fraser and Schwartz, 2012), though this conclusion is not unanimous (Chestek et al., 2011). In all cases the recording devices were floating (i.e., not rigidly fixed to the skull). This is perhaps a more suitable approach for the macaque brain, which is over an order of magnitude larger than the mouse brain. On the other hand, in a small brain like the one of a mouse, and possibly of a rat as well, our results indicate that immobile silicon probes represent a straightforward and reliable technique to obtain stable, long-term recordings of well-identified neuronal populations.

## Acknowledgements

We thank Charu Reddy and Miles Wells for technical assistance. This work was funded by Wellcome Trust Senior Investigator Awards to KDH and MC. MC holds the GlaxoSmithKline / Fight for Sight Chair in Visual Neuroscience.

